# Recontextualizing Eukaryogenesis via Computational Analysis of RNA Processing in 16,449 Archaeal Genomes

**DOI:** 10.1101/2024.08.23.609445

**Authors:** Srinivasan Kannan

**Affiliations:** University of Virginia

## Abstract

The analysis of proteins related to RNA processing reveals intriguing aspects of the evolutionary transition from archaea to eukaryotes. Eukaryagenesis is the process in which the first eukaryote came into existence, with multiple hypotheses on the order of events. These hypothesis often focus on the mitochondria and cell skeletal structure, without much discussion on eukaryotic signature proteins(ESP). ESP is integral in increasing the longevity of RNA and for the increase in the variety of proteins able to be produced which ultimately increases fitness of eukaryotes. 16,449 genomes and 10 proteins were acquired and BLAST was run for each superclass for each protein. BLAST scores were compared between superclasses and analyzed. Results for proteins such as Prp9, Rex3, Histone H2A, H2B, and Histone 3 indicate that there is no substantial difference between BLAST results implying a transitional state consistent with *E*^3^ model. The results for Smd3 and Ceg1 highlight that a group of Asgardarchaeota and Diaforarchaearchaea were different to other types of archaea. These groups likely underwent similar environmental pressures giving organisms with these genes higher fitness. These early genes evolved into their eukaryotic versions, while other genes like Histone 4, Abd1, and Lsm2 may have had ancestral prototypes present across archaea. Gene prototypes likely served different purposes, but the presence of such prototypes imply that evolution of the nucleus was likely independent from the presence of the mitochondria.

## 1. INTRODUCTION

With the discovery of Asgardarchaeota, there has been an increase in research on Archaea and their relation to eukaryotes. [14] One of the most noteworthy elements of the eukaryotic cells is the information processing machinery within. Spliceosomes and epigenetics are some of the tools eukaryotic cells use to regulate transcription, in contrast to the primary usage of operons in prokaryotic cells. Yet, the close relationship that eukaryotes have with Asgardarchaeota implies that archaea potentially have more complex tools for regulating transcription than bacteria. [14]

Archaea phylogeny is usually divided two main clusters. Cluster 1 includes the superclasses Methanomada, Acherontia, Stygia, and Proteoarchaeota, while Cluster 2 includes the two superclasses Methanotecta and Diaforarchaea [3]. Of these super-classes, Proteoarchaeota is the class that contains eukaryotes under the two-domain hypothesis [3]. While DPANN is also an important group of archaea, their placement on the phylogeny is difficult to determine [3]. Due to its unknown placement, DPANN will be exempt from the experiment. The phylogenetic relations of the superclasses of Cluster 1 and Cluster 2 are present in Figure 1.

**Figure 1.**
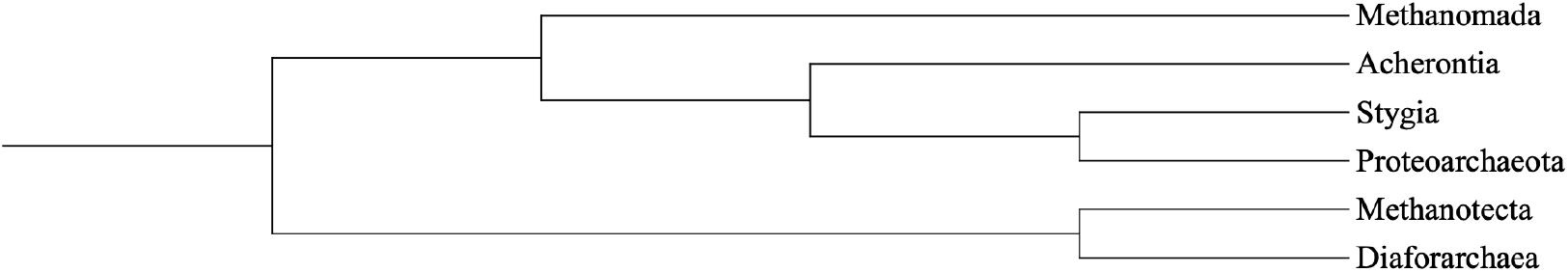
Summary of superclasses within Archaea. Diagram was created from the results in Aouad et al. [3]

Numerous protein groups are involved in transcription regulation and RNA processing. Within eukaryotes, the entire classification of transcription factors are proteins involved in regulating transcription. In addition to regulating transcription, splicing is a critical process, determining the final protein produced. Capping the mRNA ends is also a significant process as it increases the longevity of the mRNA.

Despite the role of RNA processing in eukaryotes, most models of eukaryagenesis focus on either the actin cell structure or the presence of the mitochondria [5]. Donoghue et al provides a strong analysis on the different models of eukaryagenisis and the general definition of the term [5]. The main models that will be discussed are the *E*^3^ model and the phagocytosing archaeon model (PhAT). The *E*^3^ model was proposed in Imachi et al, with the first cultivated Asgardarchaeota [9]. The study puts forward the idea that there was symbiosis between an Asgardarchaeota and other bacteria, with the bacteria entering Asgardarchaeota and becoming a Mitochondria[9]. The PhAT theory postulates that a heterotrophic archaea, which already contains a nucleus, engulfed an ancestral mitochondria. Rather than consuming the bacterium, it pursued an endosymbiotic relationship with it leading it to transform into the mitochondria. [5].

This paper seeks to analyze the sequence of events by analyzing RNA processing proteins. When analyzing current endosymbiotic models, the mitochondria and cytoskeleton are the most discussed elements. Fundamentally, these two elements are some of the biggest difference between eukaryotes and prokaryotes. Other mechanisms that are present in eukaryotes and not prokaryotes, are not often discussed, especially the presence of eukaryotic signal proteins (ESPs). The discussion of ESPs is limited because most ESP exist in all eukaryotes, but are not found in other species. However, the presence of some proteins originally classified as ESP in Asgardarchaeota provides another mechanism to determine chronology of events. With access to a wide range of Asgardarchaeota genomes now available, analysis can be done to see potential convergence of traits. Convergence of traits in multiple lineages would point to an environmental pressure selecting for eukaryagenesis.

Throughout the paper there will be discussion of environmental factors which potentially increased fitness of certain traits. Studies have estimated eukaryagenesis close to the Great Oxidation Event (GOE) [11]. Oxygen presence in the enviorment would likely have led a massive shift in the change of the adaptations of archaea, as previously existing adaptations would likely not have been as useful. Images of Korarchaeota lack of the protrusions, an adaptation, found in the two cultured Lokiarchaeota species [6] [9]. Assuming the protrusions are localized to Asgardarchaeota, the protrusions could also be used as a tool to increase horizontal gene transfer. Specifically, the protrusions grant the archaea higher surface area which could then be used to reach more free floating nucleic material. If Asgardarchaeota had an easier time picking up genetic material from its enviorment, then the reasons that the split results had a higher ratio of Asgardarchaeota compared to TACK archaeota. Further tests should be done on cultured Lokiarchaeota to better understand the true purpose of the protrusions and their role in eukaryagenesis. Another method to see the role of the protrusions is attempting to date the divergence of Proteoarchaeota into TACK and Asgardarchaeota. By dating the divergence, a better correlation can be made between Asgardarchaeota and its environment can be made. It would also help in recreating the enviorment to culture them. Dating is also valuable to better comprehend the horizontal gene transfer of evolutionary pathways. Iron reduction, methanotrophy, and denitrification were major metabolic pathways that became widespread around 2.7 million years ago [11]. These metabolic processes likely had large impacts on the enviorment, thereby creating new pressures. All of these environments jointly will be referred to as ”environmental” pressures for the rest of the paper.

## 2. RESULTS

Current tools in bioinformatics, like Benchmarking Universal Single-Copy Orthologs (BUSCO) and its many dependencies, allow for the study of many aspects of a genome [18]. One of its main dependencies, Basic Local Alignment Search Tool (BLAST), can take an input sequence of nucleotides or amino acids and search for it within the genome [18]. Returning a score that compares the input sequence with different sections of the Genome, BLAST provides a solution for assessing the presence of a sequence in a genome [18].

In addition to analysis tools, over the years, NCBI has curated an extensive database of genomes and proteins. They provide different tools like Entrez and the NCBI Command Line for ease and quick access to these genomes [17]. New tools can be created by combining these tools with tools like BLAST, which can be used to test the presence of specific proteins in specific organisms. It is important to note that the NCBI database is being constantly updated [17]. Therefore, if the command tool is used to draw genomes, the number of genomes acquired will vary. 16,449 Genomes were acquired using the NCBI Dataset Command tools. There wasn’t any normalization processes taken to make sure that each phyla had the same out of genomes. By using the maximum number of genomes, the law of large numbers would leveraged to achieve a more accurate representation of the population’s genetic diversity. However, by using all genomes, there is no random selection of genomes. By not involving randomization in the selection process, a potential systemic bias could be present in the experiment. The number of genomes is representative of all the genomes available from the command line of the date of the experiment, July 21st, 2024. The date was of arbitrary selection, but contains all genomes from in the database at the time.

Figure 2 summarizes the specific phyla of organisms selected. Phyla were chosen using NCBI’s taxonomy browser. Superclasses were searched for and the representative phyla were input in. A few selected phyla lacked genomes and were removed. The number of phyla per superclass should have been normalized. Since the final analysis conducted compares super-classes, by not having the same number of phyla per class, the results for the superclass will be more accurate than the superclass that lacks the many phyla. This is most evident by the fact that Diaforarchaea has only one phylum. Thus, the results should be taken with caution, and direct generalizations should be made with caution.

**Figure 2.**
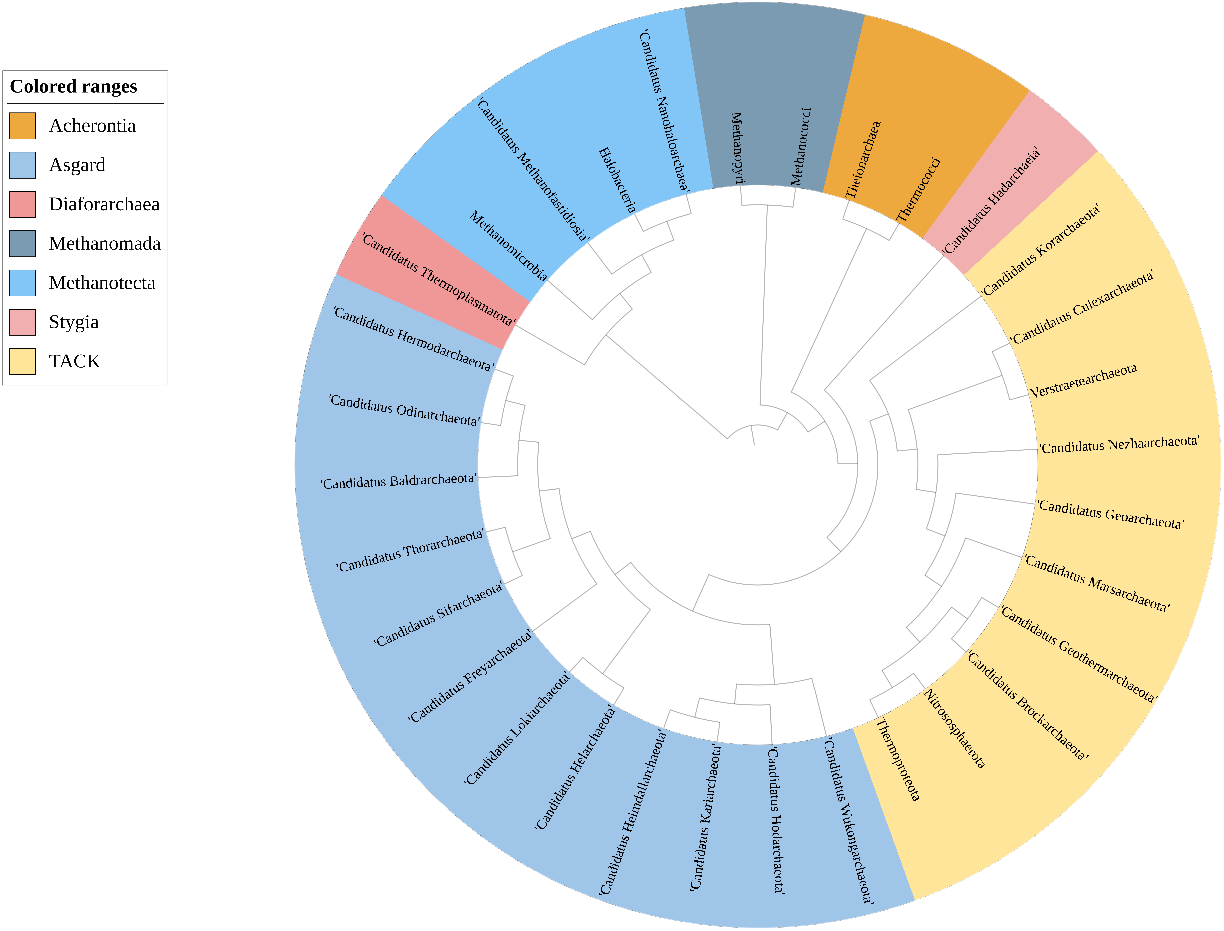
Phylogeny of all phyla used in the experiment. The tree was approximated using the results of multiple studies cited below. At the time of writing this paper, some organisms, like Freyarchaeota, don’t have specific placements on the tree. However, it can be roughly approximated using studies like Penev et al. [2] [4] [8] [10] [12] [13] [16] [19] [20]

With the genomes acquired, BLAST databases were created for each phyla. After setting up the databases, relevant protein sequences were acquired, specifically from Saccharomyces Cerevisiae. Saccharomyces Cerevisiae was selected as it was the same organism chosen by Hartman et al in their methodology of determining ESPs [7]. Despite Hartman et al being a study from 2002, its analysis of ESPs is valuable as it excluded Asgardarchaeota. By lacking relevant Asgardarchaeota analysis, the ESPs from the study can be used a basis for predicting theoretical organisms from theoretical pathways. After all, if some of the exclusive proteins is found in Asgardarchaeota and other archaea, then there is clear indications of environmental factors at play.

Then, the translated BLAST algorithm (tBLASTn) was run on all phyla. tBLASTn was selected as the appropriate BLAST algorithm due to searching a nucleic acid genome for an amino acid sequence. For high-scoring pairs (HSP) with BIT scores above 40, the respective BLAST scores were collected and used for ANOVA analysis. The BIT score of 40 is different from the 55 used in Hartman et al. [7] When assessing orthologous genes that potentially have led to different descendent proteins, it’s potentially worth decreasing the threshold value to have a greater insight on evolutionary pathways. However, conclusions should be understood with caution.

The ANOVA analysis was conducted with a significance level of 0.05 along with Turkey HSD’s pairwise comparisons were done between every phyla. Thus, per protein, a total of 528 comparisons were made. Each comparisons resulted in either a true if there was a statistically significant difference or a false if there was not a statistically significant difference. For further analysis, the comparisons were categorized by the two superclasses, the six seen in Figure 1, that were being compared. After categorization, a ratio was calculated between the number of statistically significant comparisons between the two super classes and the total number of comparisons between them. The summary of these results is graphed and can be in Figure 3 through Figure 8.

**Figure 3.**
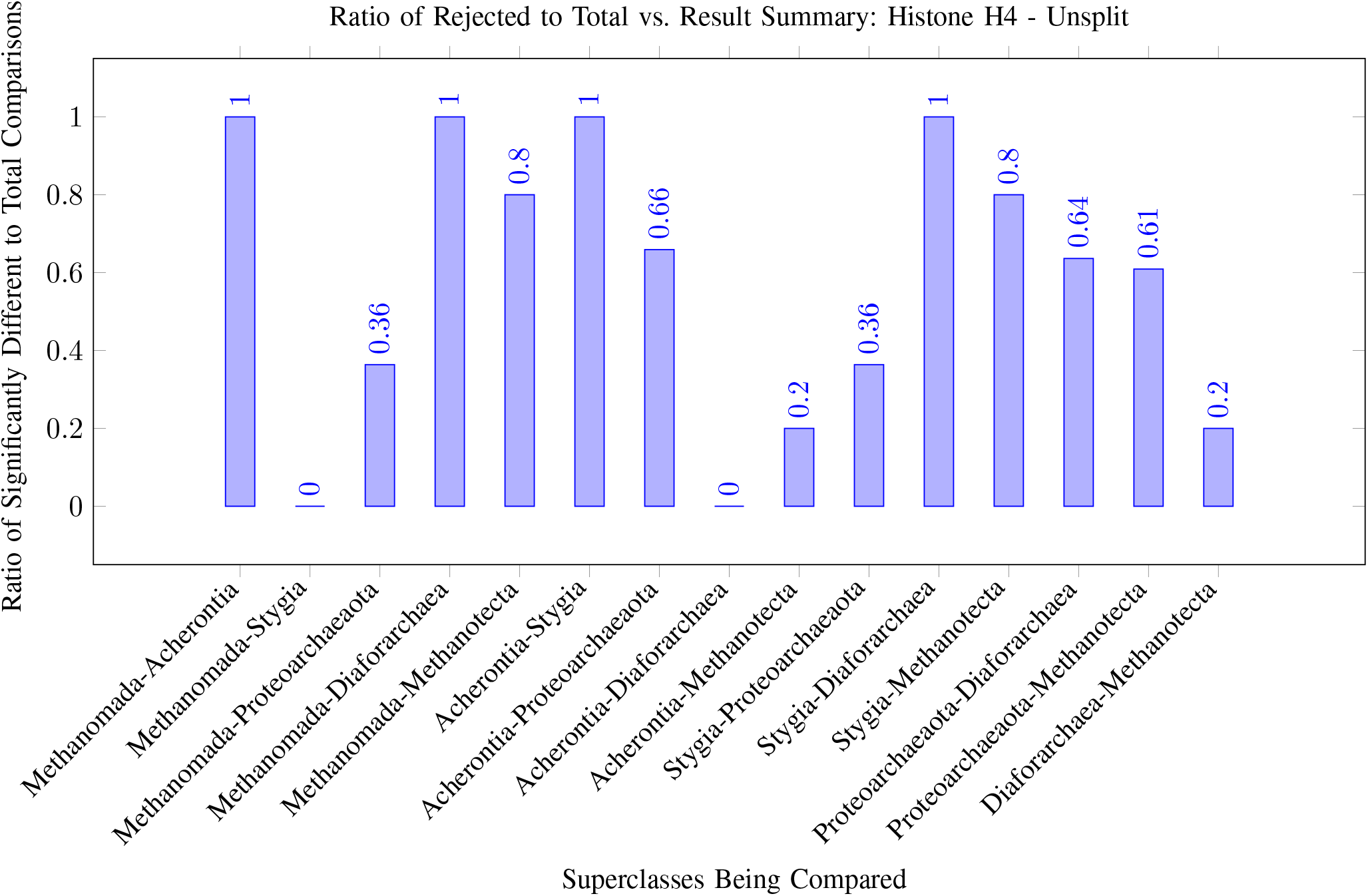
Results for Histone 4 with Proteoarchaeota

Additionally, Proteoarchaeota was split into the super phyla of TACK and Asgardarchaeota, and the ratios were recalculated and graphed. The results of the latter are in Figure 9 through Figure 14. The separation of Proteoarchaeota is to analyze the change into eukaryotes since eukaryotes are more related to Asgardarchaeota than TACK.

Supplementary Resource 1 contains details of all organisms and proteins used. It also contains the raw ANOVA results along with a copy of the graphs in Figures 3 through 14. The main difficulty in recreating the procedure is the computational space taken up by the project. Certain databases can go beyond 1 gb of space. The code used is available for use in Supplementary Resource 2. Due to amount of space that the databases take place, only the relevant code is present. The code is split into two files: one file for conducting the experiment and one files for analysis. The file for conducting the experiment takes in a list of organisms, proteins, and the significant level and outputs the result as a txt file. The file for analysis takes the result for one protein in an excel file and calculates the ratio. Since the result of the first file is in a txt file, the file should be first converted into an excel file before the use of the second file. The ratio results would be written into the excel file.

## 3. DISCUSSION

### 3.1. Smd3

Smd3 is a protein involved in splicing. Specifically, UniProt defines the role of Smd3 as a “core component” of other small nuclear ribonucleoproteins (snRNPs) like U1, U2, U4, and U5. Together, these snRNP with small nuclear RNAs (snRNA) form the spliceosome [1]. The spliceosome then serves the purpose of removing introns from a transcribed mRNA. Consequently, the protein is critical in most eukaryotes, as the lack of their presence would lead to incorrect proteins.

Figures 4 and Figure 9 imply that many changes in the snRNP gene in phyla within the Proteoarchaeota. Thus, the Smd3 gene’s ancestral version likely appeared around the emergence of early Proteoarchaeotas. With a ratio of 0.5 of statistically significant comparisons compared to a total number of comparisons, half of the tested phyla in the Proteoarchaeota superclass seem to have a gene different from the potential ancestral gene in other archaea. Examining the two superphyla in the class indicates that the TACK group has a 0.3 ratio while the Asgardarchaeota group has a ratio of 0.667. Natural selection factors could have favored the presence of the modified Smd3 gene in an enviorment that contained the ancestor of different groups of Asgardarchaeota and a few TACK archaea explaining the difference in ratios. Another possibility for the ratio discrepancy is that the protrusions, discussed earlier, allowed different Asgardarchaeota to gain the gene faster through horizontal gene transfer. By not having the same adapations that Asgardarchaeota, different species of TACK archaea were not able to gain the gene from the enviorment.

**Figure 4.**
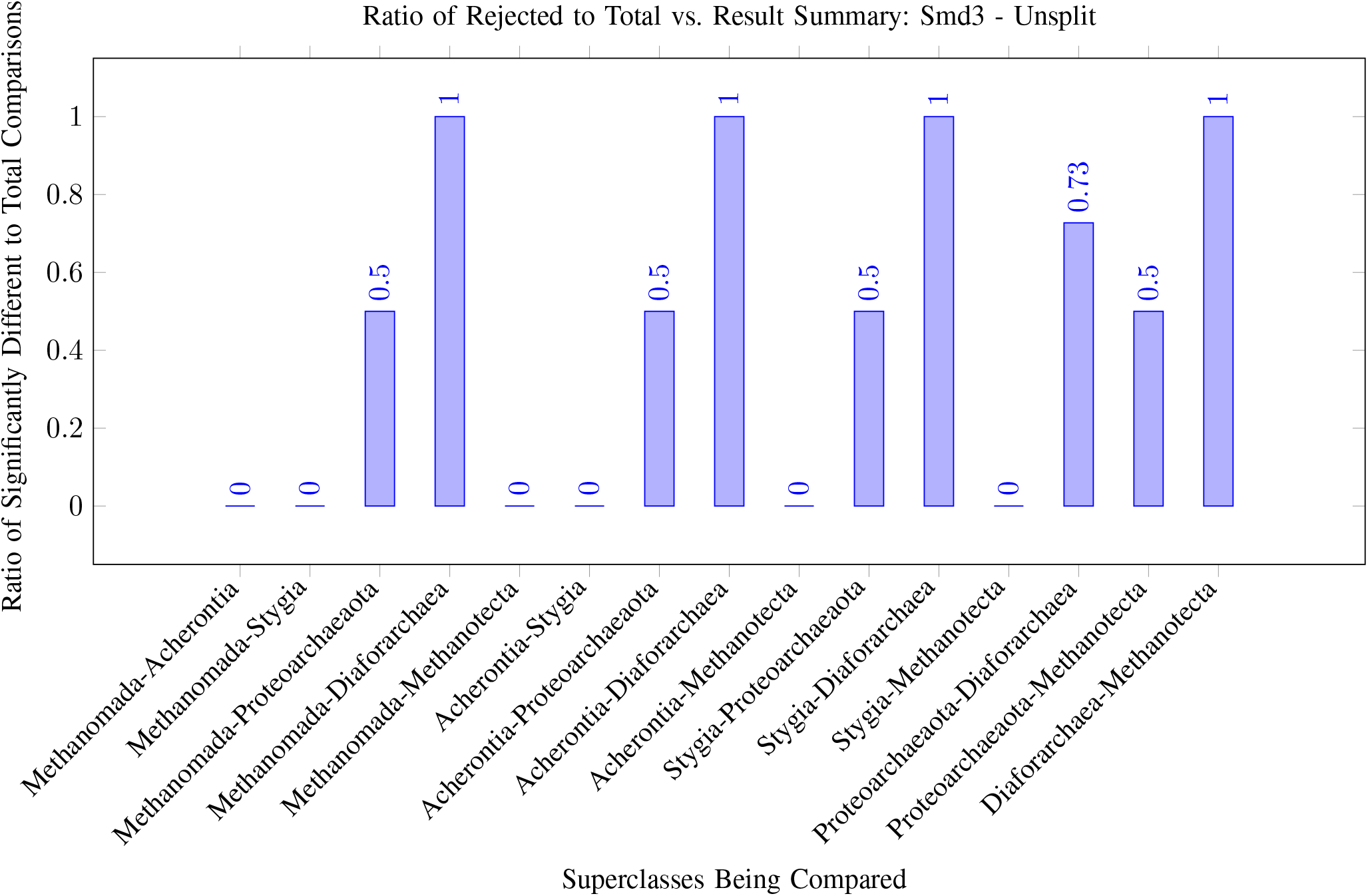
Results for Smd3 with Proteoarchaeota

TACK archaea likely gained the prototype of the gene from natural selection. The results of the analysis of the tree constructed in Figure 3 indicate there is a variety in the TACK Archaea that were significant (Culex-, Verstraete-Thermoproteota, and Nitrososphaerota) with some which are deeper within the TACK subtree implying that they are closer to other TACK Archaea than other archaea. When looking through the raw data, the difference between Verstraetearchaeota and other TACK Archaea, like Geoarchaea, leads to the conclusion that this potentially was a later change than an ancestral state. Thus, the Proteoarchaeota ancestor likely had did not have a specific ancestral gene that was the prototype to the Smd3 gene.

Diaforarchaea also has a ratio of significance compared to other archaea. Perhaps potential similar environmental pressures weakened natural preservation factors of the Smd3 gene sequence in other archaea. Despite the ratio being a 1, due to only a singular phyla being used, it is dangerous to generalize and state that the ancestor of all Diaforarchaea had the gene. In addition, the ratio between Diaforarchaea and Proteoarchaeota was 0.73, indicating that the studied Diaforarchaea phylum was different from majority of the studied Proteoarchaeota phyla. Thus, the changes that occured in the gene seem to be highly speciifc for the phyla that is similar in only a few other archaea phyla.

### 3.2. Lsm2

Lsm2 is a highly involved protein. Together with Lsm8, it forms a Lsm2-Lsm8 protein complex, which is involved in binding to the U6 snRNP. Through the binding to the snRNP, the complex is involved in splicing. Uniprot stresses the importance of the protein by stating that ”the processing of pre-tRNAs, pre-rRNAs, and U3 snoRNA” requires Lsm2. Through their statement, Uniprot highlights the importance of the protein in RNA processing. [1]

The results of the experiments, seen in Figure 5 and 10, seem to indicate a few results of note. Firstly, the dispersed results imply, unlike Smd3, a widespread form of the proto-Lsm2 gene throughout archaea. Methanomada is especially interesting as it is isolated from almost all other types of archaea. A look at the raw result indicates the two phyla of Methanococci and Methanopyri had important significant differences that led to a 0.5 ratio spread throughout the different types of archaea. For example, Methanococci did not have a statistically significant difference in terms of BLAST results compared to Diaforarchaea, while Mehtnaopyri did. Consequently, it can be stated that the 0.5 ratio could likely be due to coincidence. However, another potential explanation is that Methanomada likely had an easier time modifying the gene through the horizontal gene transfer of transformation. It would explain how the general ratio remains consistent compared to other groups of archaea.

**Figure 5.**
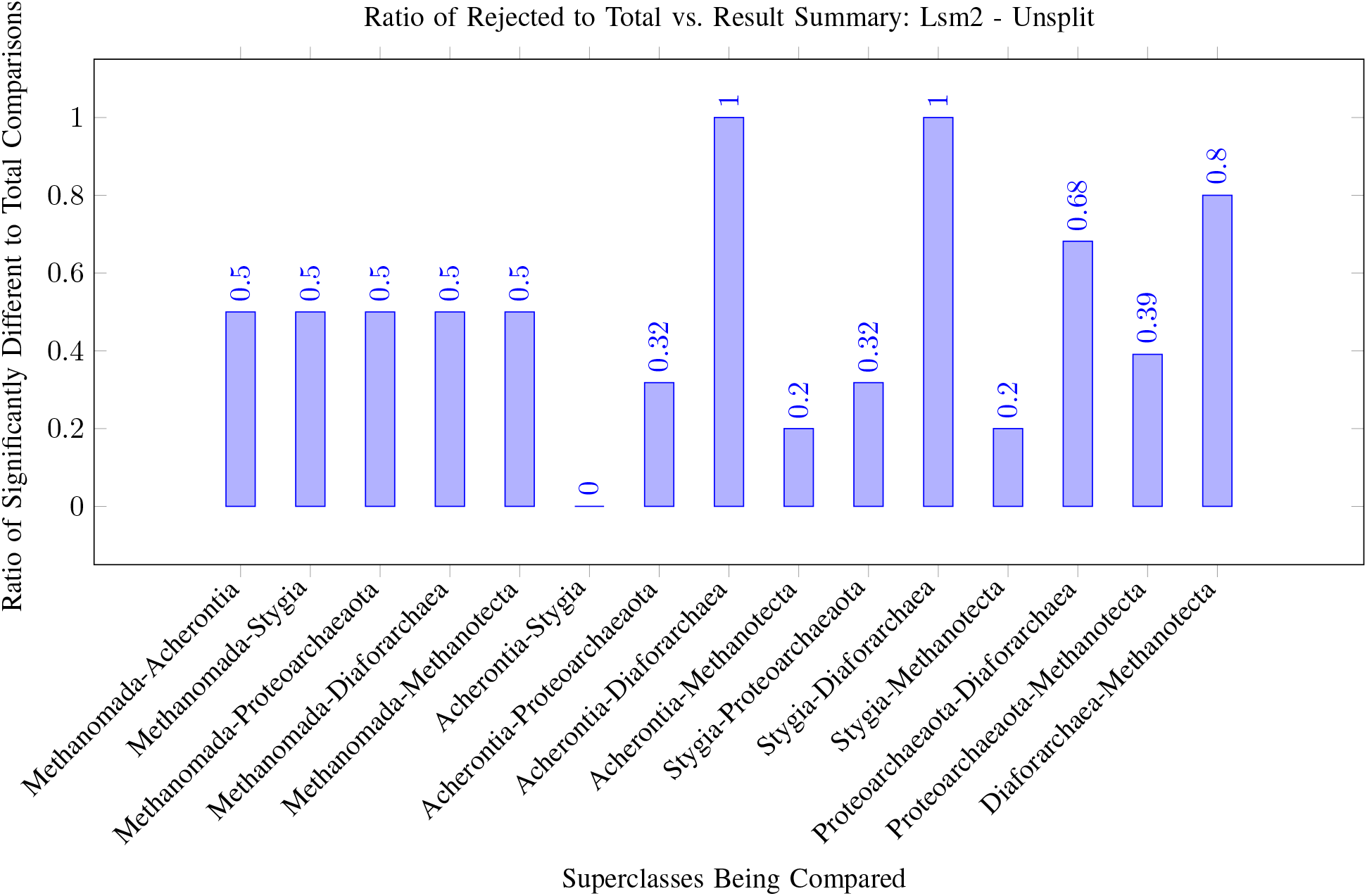
Results for Lsm2 with Proteoarchaeota

Besides Methanomada, The three main varied results came from Proteoarchaeota, Methanotecta, and Diaforarchaea. Diaforarchaea had substantially different results when compared with the other superclasses. The closest clade to Diaforarchaea also had some significant results, but the ratio of different results in Methanotecta compared to Diaforarchaea indicates that majority Methanotecta phyla were significantly different from the Diaforarchaea phyla. Due to many Methanotecta phyla being different from Diaforarchaea, the ancestor of Cluster 2 archaea did not have a common proto-Lsm2 gene.

The results of Proteoarchaeota are not too surprising. With the results of ratio between Stygia and Acherontia being 0, the change occured within the Proteoarchaeota superclass. The presence of the critical protein in eukaryotes implied that there would be more organisms that are significantly different compared to organisms that lacked it. However, it is worth noting that the difference between Asgardarchaeota and TACK is slim. Both have roughly a 0.3 ratio difference in the ratio of organisms that contain statistically significant results. Therefore, it is more likely that specific regional environmental factors increased fitness of having a proto-Lsm2 gene. It is less likely that horizontal gene transfer is involved due to the similarity between the TACK and Asgardarchaeota. If the protrusion was involved in helping horizontal gene transfer, then the lack of a high ratio of Asgardarchaeota phyla being significantly different indicates that it had a lesser role.

### 3.3. Ceg1

Ceg1 is a relatively common ESP that caps the 5’ end of the mRNA sequences. Uniprot describes it as one of the first proteins involved in capping. Capping is a cellular process that is evolutionarily conserved in eukaryotes as it helps increase the lifespan of mRNA sequences. The increased lifespan is valuable as the mRNA has to travel out of the nucleus for eukaryotes. According to Ramanathan, the capping process is a well-conserved aspect of eukaryotes. Therefore, understanding the evolution of proteins that extend mRNA longevity is necessary to understand the evolution of the nucleus. [1]

Ceg1 had similar results to Smd3. As seen in Figures 6 and 11, the groups that had significantly different results compared to others were Diaforarchaea and Proteoarchaeota. Diaforarchaea had a higher ratio of significant phyla to the total comparisons. Again, it is likely because only one phylum was studied. Figure 11, similar to Smd3, implies that environmental factors likely led to the changes of a similar number of phyla in the ancestral gene. The ratios of TACK and Asgardarcheaota had the same results throughout the different comparisons, 0.1 and 0.084. The closeness of the results imply that environmental factors had a more prominent factor like described with Lsm2.

**Figure 6.**
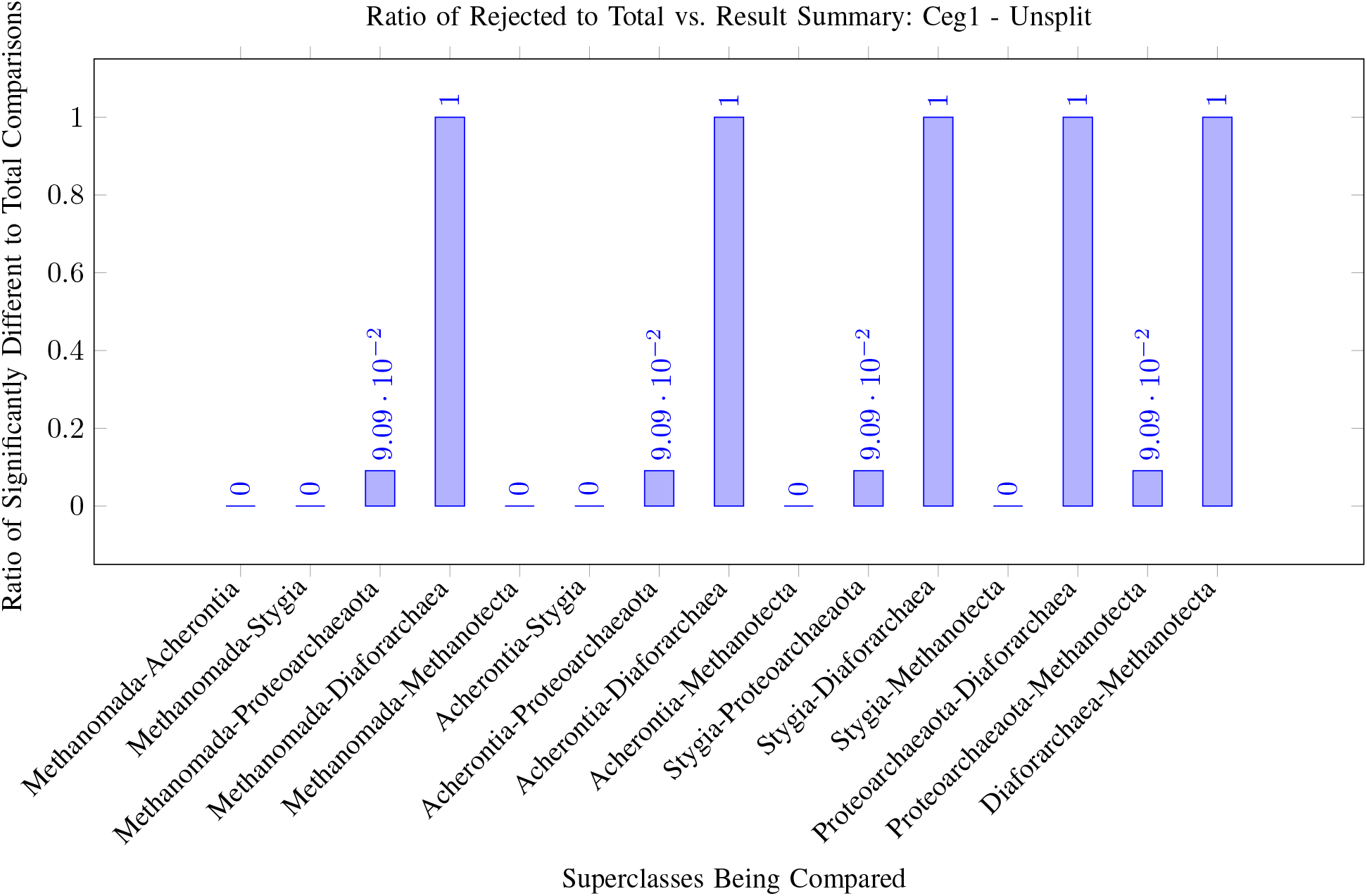
Results for Ceg1 with Proteoarchaeota

Close examination at the phyla being different, seen in Supplementary 1, indicate that Lokiarchaeota is the major Asgardarcheaota phyla with differing with other archaea phyla. Thus, the specific change into Ceg1 occured closer to Eukaryagenesis than prior. In addition to Lokiarchaeota, the two main phyla that had significantly different results from other phyla of archaea are Thermoplasmatota, a Diaforachaea phylum, and Nitrososphaerota, a TACK phylum. All three of these phylum have significantly different results. The differences between these three could be from two potential factors. It could be from the threshold BIT score being 45. Due to decrease in threshold, potential false cases of significance may have been detected. Another possibility could be that different these organisms underwent different mutations in the same gene to deal with different problems. Higher fitness in the lack of the ancestral version of the gene could have likely caused a shift in the genomes for these organisms.

### 3.4. Abd1

Abd1 is another protein involved in the capping of mRNA sequences. Thus, Abd1 is a protein involved in the longevity of the mRNA sequence, allowing transportation out of the nucleus. [1]

Unlike the results of Ceg1, which had a more standardized result like Smd3, the results of Abd1 are more similar to Lsm2. As displayed in Figures 7 and 12, the higher variation of the results imply the presence of a proto-Abd1 gene found in most archaea.

**Figure 7.**
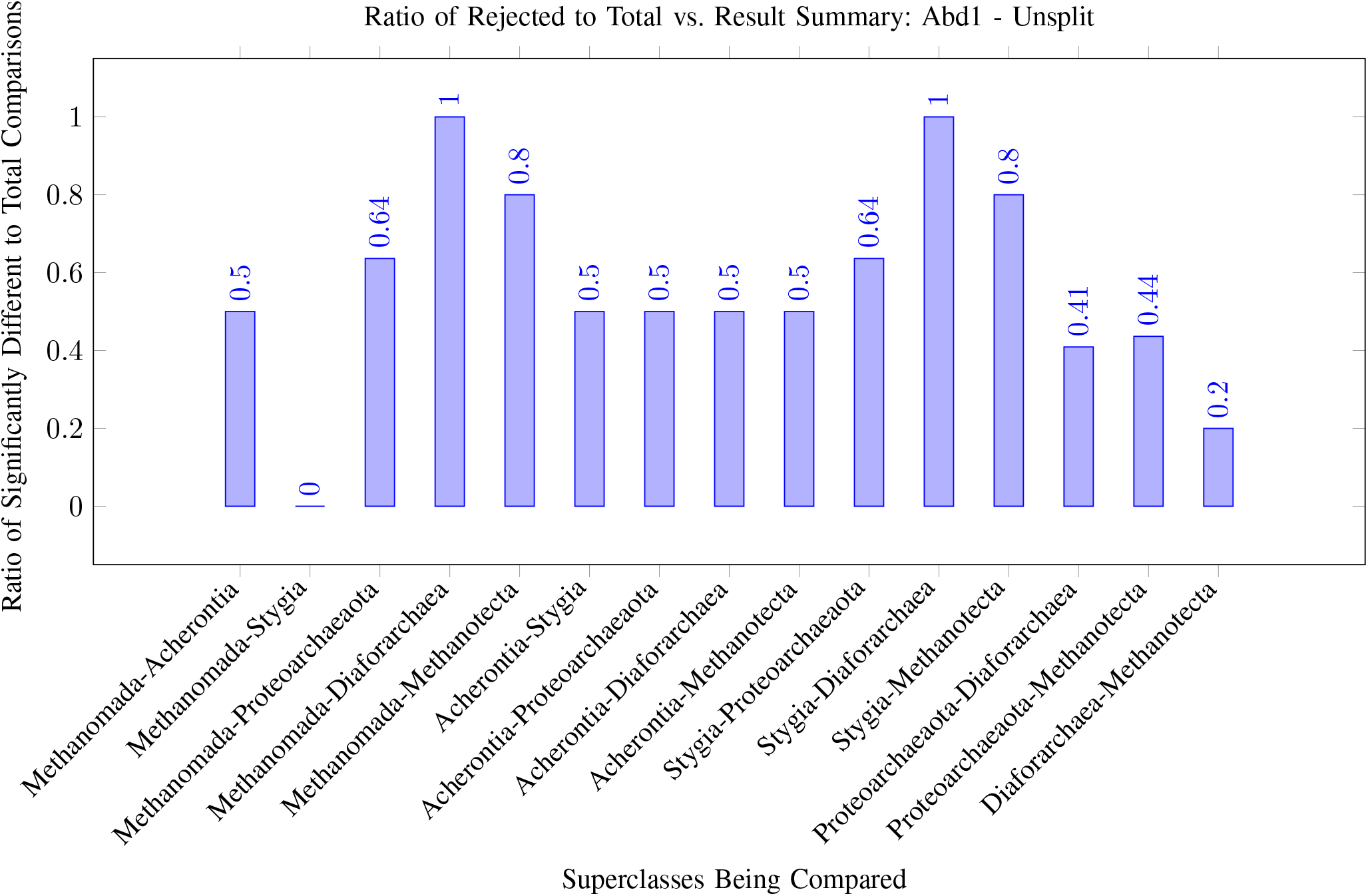
Results for Abd1 with Proteoarchaeota

**Figure 8.**
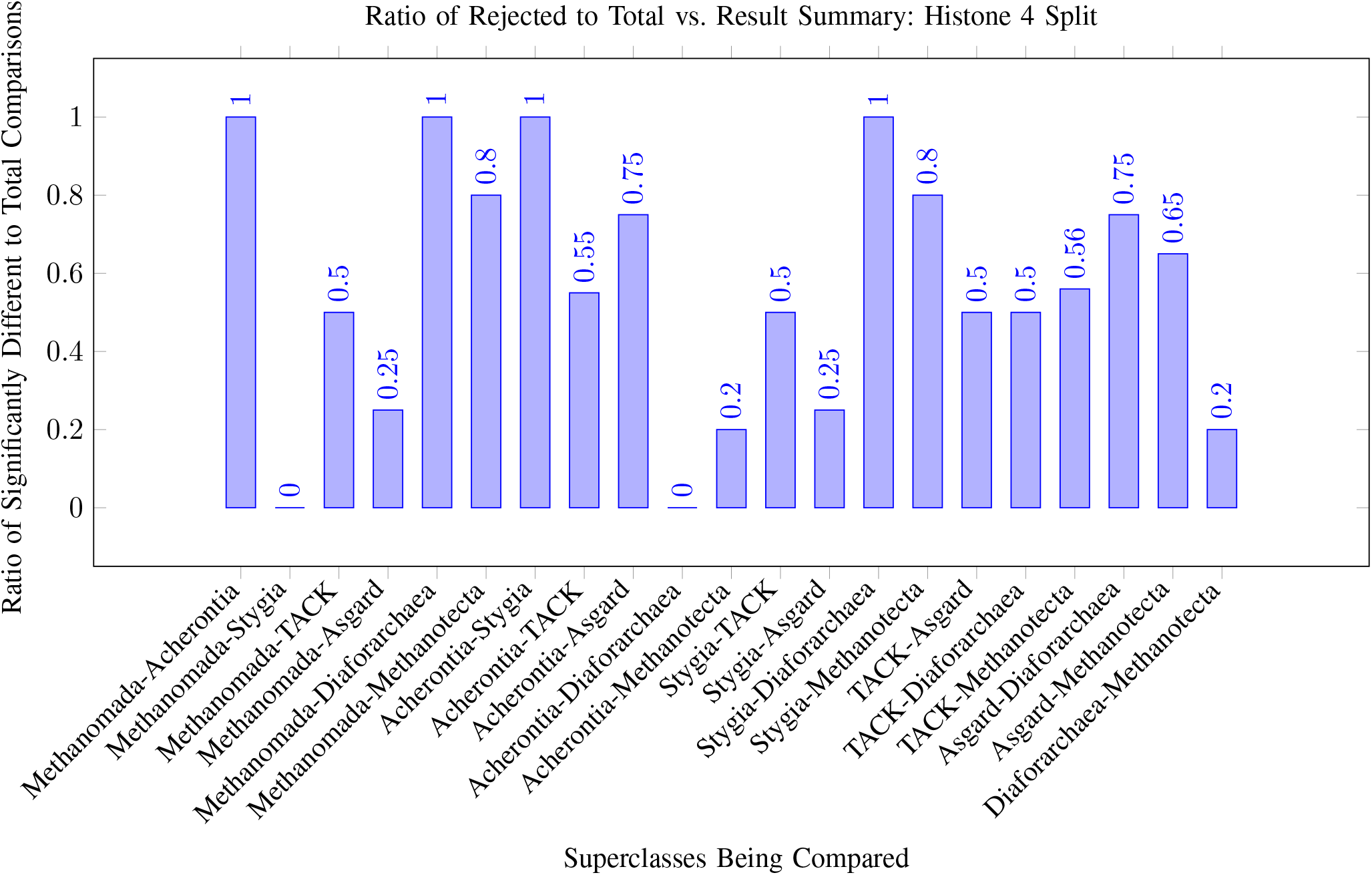
Results for Histone 4 with Proteoarchaeota split into TACK and Asgardarchaeota

**Figure 9.**
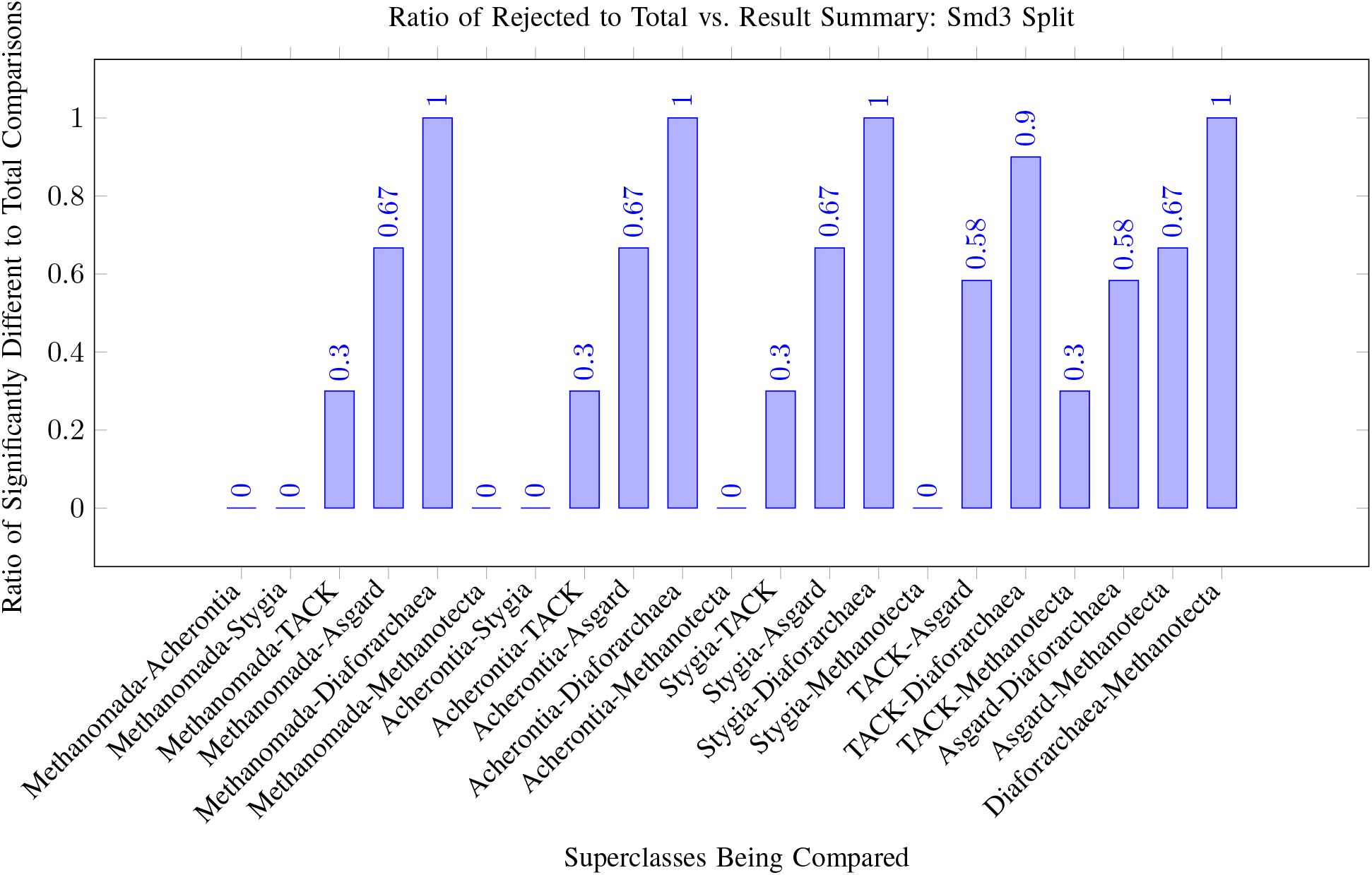
Results for Smd3 with Proteoarchaeota split into TACK and Asgardarchaeota

**Figure 10.**
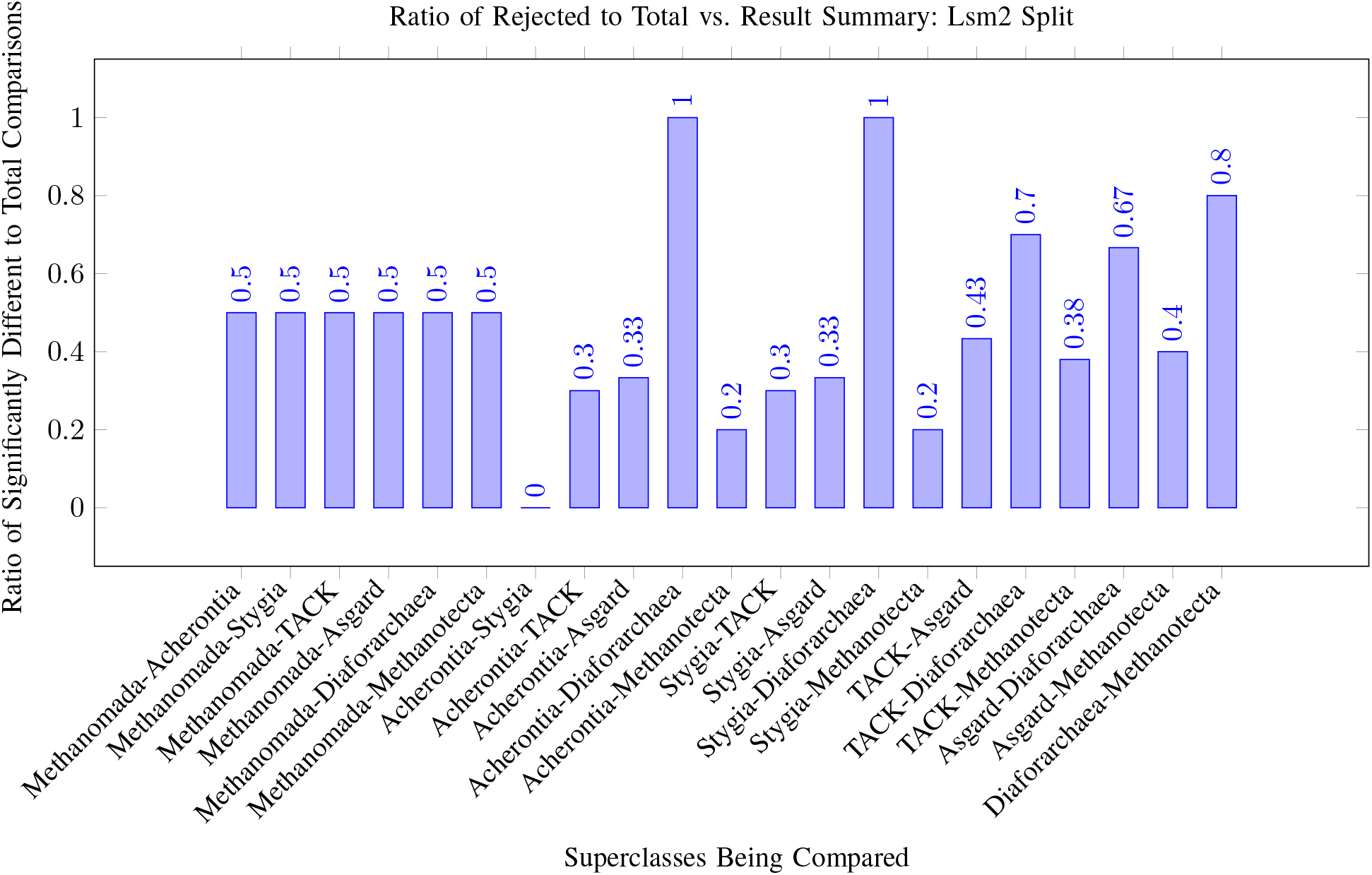
Results for Lsm2 with Proteoarchaeota split into TACK and Asgardarchaeota

**Figure 11.**
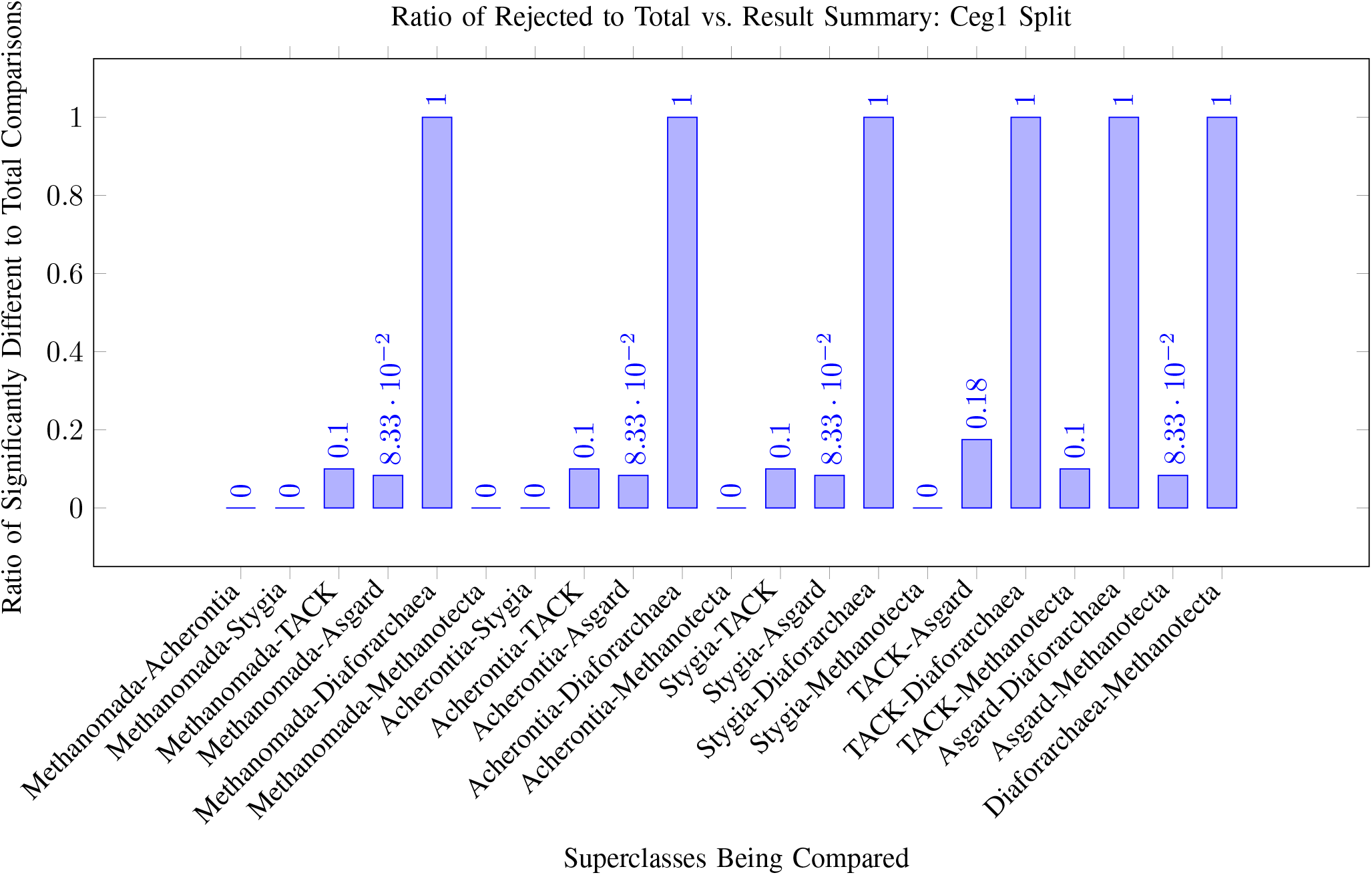
Results for Ceg1 with Proteoarchaeota split into TACK and Asgardarchaeota

**Figure 12.**
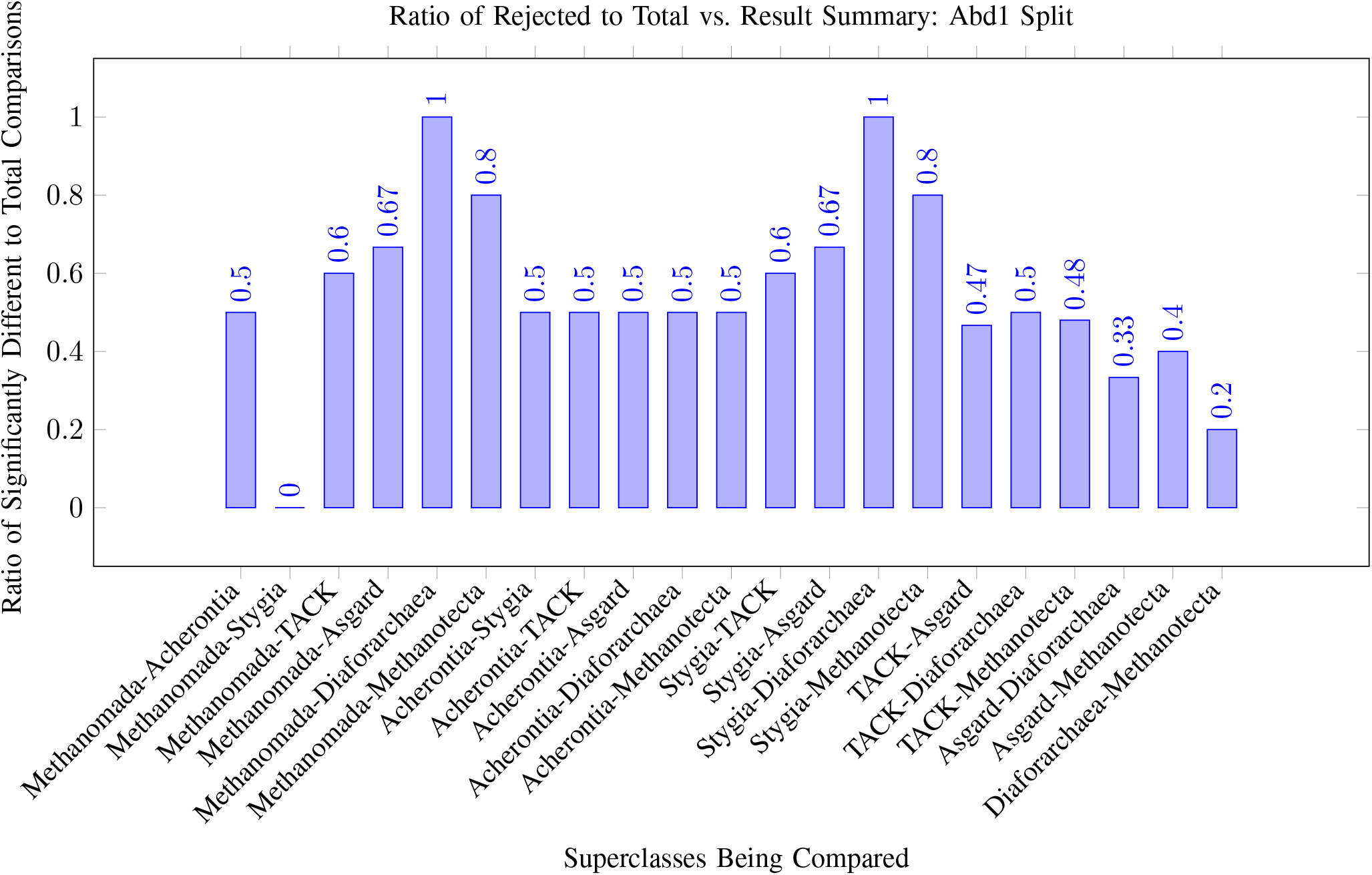
Results for Abd1 with Proteoarchaeota split into TACK and Asgardarchaeota

Acherontia has results similar to Methanomada in Smd3, having a 0.5 ratio when compared to all other superclasses of archaea. The raw results imply that the Acherontia varient of the gene was prone to change, changing depending on their environment. Since the two Acherontia phyla studied, Thermococci and Theionarchaea, are significantly different from each other, the proto-Abd1 gene in Acherontia evolved in different ways. The further change in only Acherontia is seen with in Methanomada and Stygia comparison ratio of 0. The lack of signifcant difference in the prototype of the gene implies that there was a specific change that happened in Acherontia leading it to be more susceptible for change in the environment. The 0.2 ratio between the superclasses of Methanotecta and Diaforarchaea indicates that only one phyla of the Methanotecta phyla, Methanonatronarchaeia, was different from the studied Diaforarchaea phyla. Considering that only 1 Methanotecta phyla was significantly different, it is more likely that the proto-Abd1 gene changed in this phyla alone. However, since only one Diaforarchaea was studied, the results should be taken with caution. However, an interesting note is that both Diaforarchaea and Methanotecta have high ratios when compared to the superclasses of Methanomada and Stygia. The vast difference in ratios implies a massive change in the gene during the split of Cluster 1 and Cluster 2 archaea as majority of the phyla in Methanmada and Stygia are different from the ones in Diaforarchaea and Methanotecta. Most of the Cluster 2 archaea seemingly maintaining the ancestral state with the exception of Methanonatronarchaeia.

Majority Proteoarchaeota phyla are also significantly different from the Stygia and Methanomada phyla. Considering that less Proteoarchaeota phyla are different from Diaforarchaea and Methanotecta, major changes occurred to the proto-Abd1 gene. These changes highlight a progression towards the Eukaryotic version of the Abd1 gene. Hemidallarchaeota, an archaea that is closely related to the Last Eukaryotic Common Ancestor (LECA), is significantly different from the Methanomada and Stygia phyla. From the difference between Heimdallarchaeota and the Methanmada-Stygia group, an implication that the change to the Abd1 gene would have given a higher fitness for Proteoarchaeota in the environment that their developing in. However further tests with a higher BIT threshold should be done to confirm if the Abd1 gene has a homologoue in Hemidallarchaeota.

### 3.5. Histone H4

Histone is a group of proteins involved in epigenetics and the conservation of certain genes. The presence of histones helps prevent mutations in other genes during DNA replication, but they also are involved in transcription regulation. While there are many types of Histones, Histone H4 is a specific protein that is part of the nucleosome. [1]

An initial look at the results in Figures 3 and 8 shows that there are two main groups of results: one group being Methanomada and Stygia and the other group being Acherontia and Diaforarchaea. The results between these two groups indicate no significant difference within the group, but they are significantly different from the other group (ratios of 0 and 1). While the results potentially came from the low number of phyla used, the stark differences are worth noting. Since the divergence of Acherontia occurred after Methanomada but before Stygia, it is more likely that Acherontia phyla evolved the proto-Histone 4 gene differently. If the Acherontia phyla developed in a similar enviorment as Diaforarchaea, then it is likely that horizontal gene transfer or due to convergent evolution.

The ratio difference between Diaforarchaea and Methanotecta being 0.2 implies that some Methano-tecta archaea, like Methanonatronarchaeia phyla studied, evolved differently. Environmental factors and the process of transformation likely were the driving factors of the difference. A 0.8 ratio between Methanotecta and Methanomada implies Cluster 2 archaea’s ancestral state for the proto-Histone 4 gene was likely different from Cluster 1’s proto-Histone 4 gene. Environmental factors also likely led to the change of Acherontia phyla to be more akin to the Cluster 2 archaea. It is very similar to the results of Abd1 where Methanomada and Stygia had no signficant difference while the superclass between, Acherontia, had phyla different from both. However, unlike Abd1, all the studied phyla in Acherontia were different from the phyla in Methanomada and Stygia.

The idea that the Acherontia change was separate from the rest of Cluster 1 is supported by the ratio of comparison with Proteoarchaeota phyla. By being signficantly different most Proteoarchaeota, Acherontia phyla likely developed seperately and the same environmental factors, for one reason or another, did not have substantial impact on the fitness of Proteoarchaeota phyla.

The ratio difference between the Asgardarchaeota and TACK superphyla compared to other superclasses suggests that that there was a significant impact of the environment on these superphyla. Despite having a 0.5 ratio of being different from each other, the lack of consistent ratios when compared with other superclasses indicates high variablity in the potential proto-Histone 4 gene in these organisms.

### 3.6. Prp9, Rex3, Histone H2A, Histone H2B, Histone 3, and Limitations

The experiment was conducted for all of these proteins as well. However, as seen in the raw results, there are no significantly different results. The only exception is with Histone 3, between the phyla of Korarchaeota and Methanomicrobia. The singular difference implies that there were likely changes that occurred with the ancestral gene of Histone 3, but there weren’t any factors that pushed for one over the other. Any factors that did apply were minor. The impact of minor factors likely led to the significant difference between the two phyla. However, considering the two phyla come from different clusters, implying more of an environmental effect than a genetic one.

The lack of significance in the results of the protein could imply two significant results. The lack of significant differences insinuates that the closest gene to a given protein only emerged close to or after the first eukaryotes. The other explanation would be that there is no signficant difference between the gene in any archaea. Yet, to an extent, both have their merit. The raw results for Prp9, a protein involved in the early stages of splicing, had all comparisons result in zero [1]. The raw results for Rex3, a protein involved in the later end of splicing, and the other 2 Histones had differences in ANOVA results, just none that was enough to indicate significance [1]. Since some proteins had comparison results of a 0 while others did not imply that there was likely a protogene for some proteins in some archaea, but not different enough to warrant a significant difference from other results. The lower results of the BLAST score imply that the genes likely evolved more after the first eukaryote cell formed.

A main limitation of the experiment is the lack of analysis of genomes before usage. The initial methodology for the experiment involved using BUSCO to score each genome and only let genomes of a certain threshold to be used. However, the lack of the right computer architecture resulted in skipping the BUSCO process. Despite the lack of BUSCO, it is worth considering the law of large numbers. With over 16,000 genomes being used, the results should be a good representative of the overall population.

## 4. Conclusions

Analysis of these proteins highlights a few interesting aspects of Eukaryagenesis. The lack of large statistically significant results for the proteins Prp9, Rex3, Histone H2A, Histone H2B, and Histone 3 implies a possible transition state from which archaea evolved into eukaryotes.

The prototype genes of these proteins may or may not have the same purpose. Studies have shown that certain genes used for one mechanism may not be used in another mechanism, as demonstrated by the origin of the flagella. Furthermore, while the rate of single mutations is small, it should be understood with the context of the number of archaea and generations passed. Therefore, direct homologous genes may not portray the full scope of archaeal evolution. Consequently, the threshold was decreased from 55 to 40. By standardizing everything else, the scores show clear trends that improves on existing knowledge.

Eukaryotes are defined as cells that contain a nucleus, yet these results seem to push the idea that the cell first gained the Mitochondria. Horizontal gene transfer mitochondria can explain the presence of these different proteins that aren’t present in the genomes of archaea. Proteins like Smd3 and Ceg1 likely had earlier forms emerging before the first eukaryotes showed signs of developing in close promixity within lineages in Proteoarchaeota. The emergence of these genes was likely due to a combination of environmental factors and higher fitness in those that changed to better match the environment. These prototype genes would then evolve into their respective version in eukaryotes. In contrast, genes like Histone 4, Abd1, and Lsm2 likely had an ancestral prototype in all archaea which later evolved into their eukaryotic version due to not having one or two specific superclasses with clear distinct results compared to others.

The study carries two main implications. The classification of eukaryotes have usually been with the presence of the nucleus. At the time of writing this study, all found eukaryotes have a nucleus and some form of a mitochondria. However, by lacking ESP in the closest archaeal phylums, the possibility of rare eukaryotes in a transition state should be considered. The results of the Smd3 and Ceg1 proteins imply that environmental factors increased fitness of archaea with certain proteins. These environmental factors have likely drastically changed. Potential ”transitional eukaryotes” would be hard to find as the environmental factors that they were born from are not the same factors present today. There is a possibility that almost ”transitional eukaryotes” are now extinct due to environmental change. It is important to consider them as certain nuclear factors had to be in place before the nuclear membrane fully formed. Otherwise, the rate of mRNA degradation would have decreased the overall fitness of the first eukaryotes.

Quotes are used when discussing transitional eukaryotes as they likely are more similar to Asgardarchaeota than modern day eukaryotes. These transitional eukaryotes would be very similar in nature to the *E*^3^ model of eukaryagenesis [5]. The hypothetical organism would likely lack a strong structured nuclear membrane, or it would have some other mechanism in which the mRNA can begin translating quickly. The intermediate step with mitochondria would explain the presence of other proteins that seemingly have chaotic patterns in other eukaryotes. The development of proteins involved in splicing further suggests that splicing may have been used as a tool to produce relevant proteins for the environment. Being able to produce a variety protein as a unicellular organism would yield higher fitness in those species. Lack of these elements would lead to competing proto-eukaryotes to go extinct from resource competition. However, with the discovery of first Asgardarchaeota in the middle of 2010s, there is a possibility of finding these transitional eukaryotes in an unexpected environment [15].

## Supporting information

(Already in PDF as an Google Spreadsheet Link) Supplementary Resource 1: Raw Results

## 5. Appendices

Supplementary Material 1 contains all collected data. Supplementary Material 2 contains all used code. For specific BLAST databases used, please contact through email.

[Supplementary Material 1: Raw Results](https://docs.google.com/spreadsheets/d/12c8hrwaz-wqr092ceDNKNLN-4i25lzARRhypuW_YuZU/edit?gid=560610447#gid=560610447)

[Supplementary Material 2: Code](https://github.com/SaiSri-bit/Genome-Statistical-Analysis)

